## Supplemental Information for "Dynamic ubiquitination determines transcriptional activity of the plant immune coactivator NPR1"

**Tables S1 – S3**

**Figures S1 – S5**

**Table S1. Related to Figure 3. SA-induced genes in WT, *ube4* and *npr1* plants determined by RNA-Seq.**
See supplemental file.

**Table S2. List of oligonucleotides used.**

| PCR product | Purpose | Sequences (5'-3') |
|---|---|---|
| *PR1* | qPCR | F - CTAAGGGTTCACAACCAGGC<br>R - AAGGCCCACCAGAGTGTATG |
| *WRKY18* | qPCR | F - AGAAGGTACAACGCAGCGCAGA<br>R - TGCGTCCCTTCGTATGTCGCTACA |
| *WRKY38* | qPCR | F - CCGGTTTACCGAACCACTTA<br>R - GGCTTTCCTTCTCCTGATCC |
| *WRKY62* | qPCR | F - GCCTACACCAAGGACCAGAA<br>R - AGAGGTGGAGGAGGAGAAGC |
| *NIMIN1* | qPCR | F - CACGGAAACGTAGACGAGAA<br>R - CCCGTACGACACTGAGAGAA |
| *BIP1* | qPCR | F - CCACCGGCCCCAAGAG<br>R - GGCGTCCACTTCGAATGTG |
| *PR5* | qPCR | F - ACTGTGGCGGTCTAAG<br>R – CGTGGGAGGACAAGTTT |
| *TRXh5* | qPCR | F - CATACCCTCGAAGTTTGGAACGAGA<br>R - TTGCCTCAACTTTGAATTCCTGAGC |
| *PR2* | qPCR | F - CAGATTCCGGTACATCAACG<br>R - AGTGGTGGTGTCAGTGGCTA |
| *NPR1* | qPCR | F - CTAAAACCGTGGAACTCGGG<br>R - TCTCTTGTATTTCCATGTACCTTTGCT |
| *PR1* promoter *as1* element | ChIP qPCR | F - AGTGTATACAATGTCAATCGGTGATCTT<br>R - GCCGCCACATCTATGACGTA |
| *UBP6* | TOPO cloning | F - CACCATGCCTACAGTAAGCGTGAAG<br>R - TTACATGGAGACGAAGCGGGC |
| *UBP6(C113S)* | Site-directed mutagenesis | F - CTTGGCAACACGTCTTACATGAACTCC<br>R - GGAGTTCATGTAAGACGTGTTGCCAAG |
| *UBP6* | pET28a cloning | F - GACGAATTCATGCCTACAGTAAGCGTGAAG<br>R - ACGTGTCGACTTACATGGAGACGAAGC |
| *cul3a* | genotyping | F - CTTAACCGTTTAAAATGGGCC<br>R - GTCAGATGACGCAGAAAGGAG |
| *cul3b* | genotyping | F - GAGGGAAGACGGTGGAAATAG<br>R - AAATGCTCCTCCTTGAGCTTC |
| *ube4-2* | genotyping | F - GAACTCGTCTGGTATTTCCCC<br>R - GAGCTTGCCATGACTTTGAAC |
| *ics1/sid2-2* | genotyping | F - CAATCTTGATGCTCTGCAGCTTC<br>R - GAAGATAGTTGAACCAAGG |
| *npr1-1* | genotyping (CAPS) | F – CTCGAATGTACATAAGGC<br>R - CGGTTCTACCTTCCAAAG |
| *uch3-1* | genotyping | F - CGAATCTGATTTTGTGATTCG<br>R - GAATTTGGTAGGTGCATAGCG |
| *ubp6-1* | genotyping | F – TGGTCCAAGTGGATGGATAAG<br>R - TGCAAATGGAAGTGGAGAATC |
| *ubp7-1* | genotyping | F – CCCATACTTATGGTGCATCATC<br>R - AGGCGACAAATATCCAGGTTC |

### Table S3. List of reagents and materials used

| Reagent or material | Source or reference | Identifier |
|---|---|---|
| Antibodies | | |
| Mouse monoclonal anti-GFP | Roche | Cat# 11814460001 |
| Rabbit polyclonal anti-S5a | Abcam | Cat# ab60101 |
| Rabbit polyclonal anti-NPR1 | This paper | N/A |
| Rabbit polyclonal anti-GAPDH | Sigma-Aldrich | Cat# G9545 |
| Rabbit polyclonal anti-pS11/15 NPR1 | (Spoel et al., 2009) | N/A |
| Rabbit polyclonal anti-GFP (ChIP grade) | Abcam | Cat# ab290 |
| Mouse monoclonal anti-Ubiquitin (FK2) | Millipore | Cat# 04-263 |
| Mouse monoclonal anti-FLAG M2 affinity gel | Sigma-Aldrich | Cat# A2220 |
| Rabbit monoclonal anti-FLAG | Sigma-Aldrich | Cat# F7425 |
| Rabbit polyclonal anti-RPN6 | Upstate | Cat# 09-281 |
| Mouse monoclonal anti-Ubiquitin (P4D1) | Santa Cruz Biotechnology | Cat# sc-8017 |
| Mouse monoclonal anti-HA | ThermoFisher | Cat# 26183 |
| Mouse monoclonal anti-T7 | Millipore | Cat# 69522 |
| Bacterial and Virus Strains | | |
| *Escherichia coli* strain TOP10 | Invitrogen | Cat# C404010 |
| *Escherichia coli* strain BL21(DE3) | Spoel lab stock | N/A |
| *Agrobacterium tumifaciens* strain GV3101 (pMP90) | Spoel lab stock | N/A |
| *Pseudomonas syringae* pv *maculicola* ES4326 | Spoel lab stock | N/A |
| Chemicals, Peptides, and Recombinant Proteins | | |
| PR-619 | Abcam | Cat# ab144641 |
| NSC632839 | Abcam | Cat# ab144599 |
| WP1130 | Cayman Chemical | Cat# 15227 |
| P2207 | LifeSensors | Cat# SI9699 |
| TCID | LifeSensors | Cat# SI9679 |
| MG132 | Cayman Chemical | Cat# 10012628 |
| USP2 Catalytic Domain | Boston Biochem | Cat# E-504 |
| 26S Proteasome (Ub-VS treated) | Ubiquigent | Cat# 65-1020-010 |
| Poly-ubiquitin (Ub3-7) K48-linked | Boston Biochem | Cat# UC-220 |
| Poly-ubiquitin (Ub3-7) K63-linked | Boston Biochem | Cat# UC-320 |
| Di-ubiquitin K48-linked | Boston Biochem | Cat# UC-200B |
| Di-ubiquitin K63-linked | Boston Biochem | Cat# UC-300B |
| HA-Ubiquitin-Vinyl sulfone | Boston Biochem | Cat# U-212 |
| Critical Commercial Assays | | |
| SuperScript II | Invitrogen | Cat# 18064014 |
| QuikChange Site-Directed Mutagenesis Kit | Agilent | Cat# 200519 |
| GFP-Trap A | Chromotek | Cat# gta-20 |
| Deposited Data | | |
| RNA-Seq | Array Express at EMBL-EBI | E-MTAB-7369 |
| Experimental Models: Organisms/Strains | | |
| *Arabidopsis*: *cul3a cul3b* | (Spoel et al., 2009) | SALK_046638 SALK_098014 |
| *Arabidopsis*: *ics1/sid2-2* | (Wildermuth et al., 2001) | N/A |
| *Arabidopsis*: *ube4-2* | (Sessions et al., 2002) | SAIL_713_A12 |
| *Arabidopsis*: *npr1-1* | (Cao et al., 1994) | N/A |

| | | |
|---|---|---|
| *Arabidopsis*: *npr1-0* | This paper, (Alonso, 2003) | SALK_204100 |
| *Arabidopsis*: *35S::NPR1-GFP npr1-1* | (Kinkema et al., 2000) | N/A |
| *Arabidopsis*: *ubp12-2w* | (Cui et al., 2013) | GABI_742C10 |
| *Arabidopsis*: *uch3-1* | This paper, (Alonso, 2003) | SALK_140823 |
| *Arabidopsis*: *ubp6-1* | This paper, (Alonso, 2003) | SALK_108832 |
| *Arabidopsis*: *ubp7-1* | This paper, (Alonso, 2003) | SALK_014223 |
| Recombinant DNA | | |
| pENTR-D-TOPO | Invitrogen | Cat# K240020 |
| pEarleyGate 202 | ABRC (Earley et al., 2006) | Cat# CD3-688 |
| pGEX-6P-1 | GE Healthcare | Cat# 28-9546-48 |
| pET28a | Novagen | Cat# 69865 |
| Software and Algorithms | | |
| Strand NGS | Avadis® | N/A |

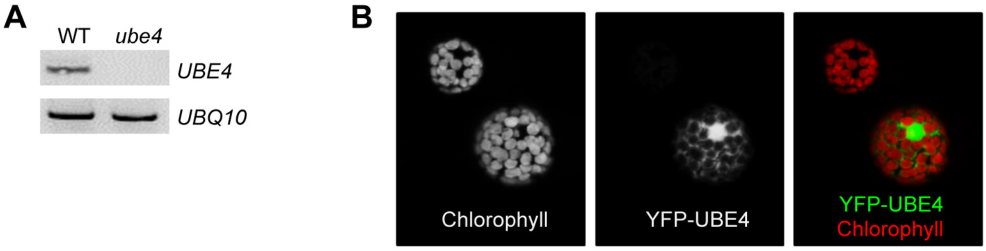


**Figure S1. Related to Figures 1 and 2. UBE4 knockout confirmation and cellular localisation.**

**(A)** Expression of *UBE4* was analysed by RT-PCR in the stated genotypes using primers specific to *UBE4* or *UBQ10* as a loading control.

**(B)** *35S::YFP-UBE4* was transformed into protoplasts and subcellular localization analysed by confocal microscopy. Left: Auto-fluorescence of protoplasts. Middle: *35S::YFP-UBE4*. Right: Merged image.

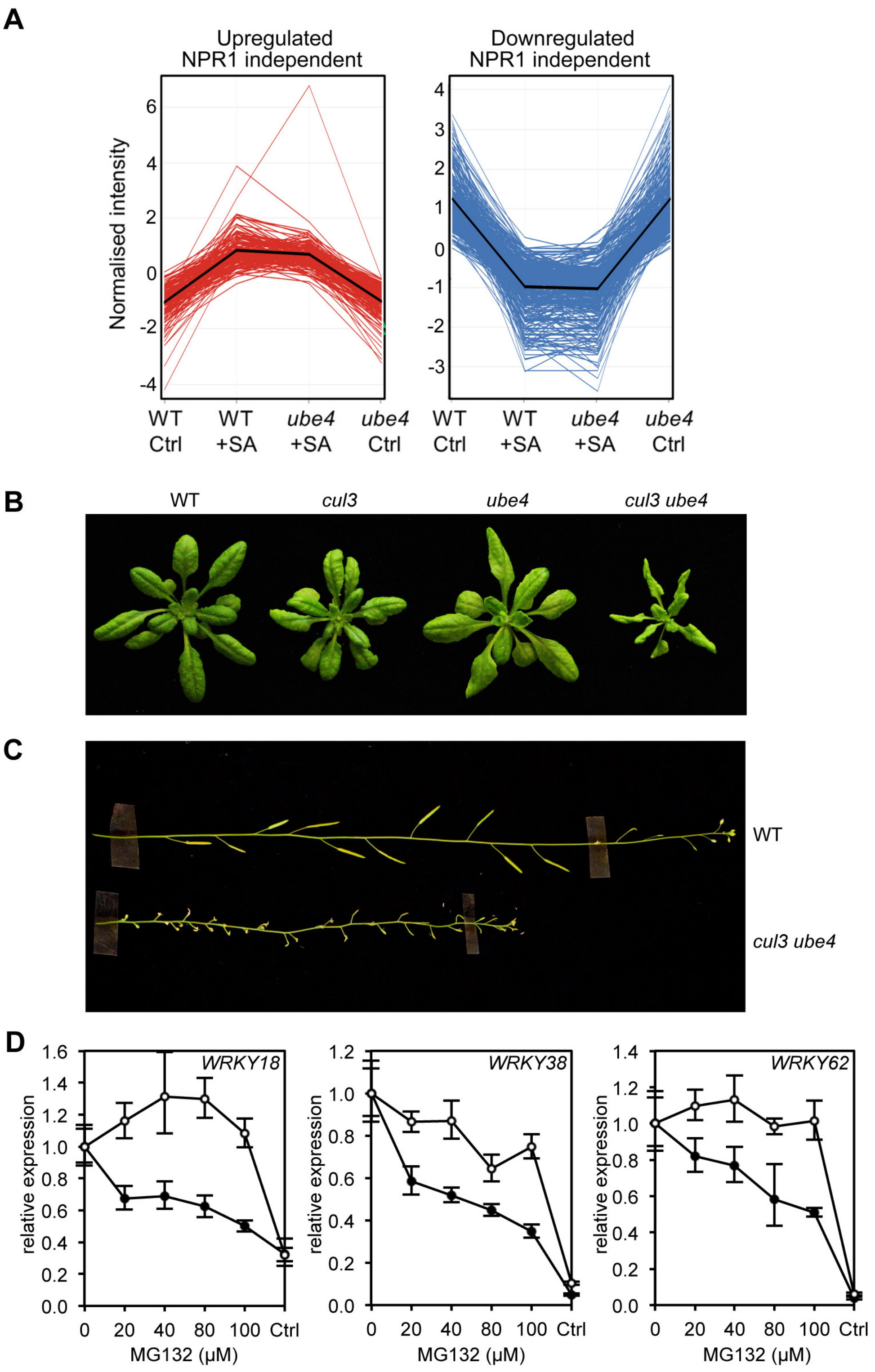


**Figure S2. Related to Figure 3. Processive ubiquitination controls transcriptional activity of NPR1**

**(A)** Expression profiles of SA-responsive, NPR1-independent genes. Seedlings treated with water (Ctrl) or 0.5 mM SA for 12h were analysed by RNA-Seq. Only genes that were induced (left) or repressed (right) ≥2-fold by SA in WT and/or *ube4* plants and showed ≤1.5-fold difference in expression in *npr1* mutants are shown (Benjamini

Hochberg FDR, 2-way ANOVA $p \leq 0.05$). Mean expression patterns are indicated by black lines.

**(B)** Morphological phenotypes of 4-week-old plants of the indicated genotypes.

**(C)** Inflorescence phenotypes of 7-week-old plants of the indicated genotypes.

**(D)** WT (closed circles) and mutant *ube4* (open circles) seedlings expressing *35S:NPR1-GFP* were treated with 0.5 mM SA for 4h followed by the addition of indicated concentrations of MG132 for a further 2h. *PR1* gene expression was determined and normalised relative to constitutively expressed *UBQ5*. MG132 treatments as well as a control (Ctrl) that received 4h of water treatment followed by the addition of vehicle (DMSO), were plotted relative to maximal SA-induced *PR1* expression. Data points represent mean ± SD (n=3).

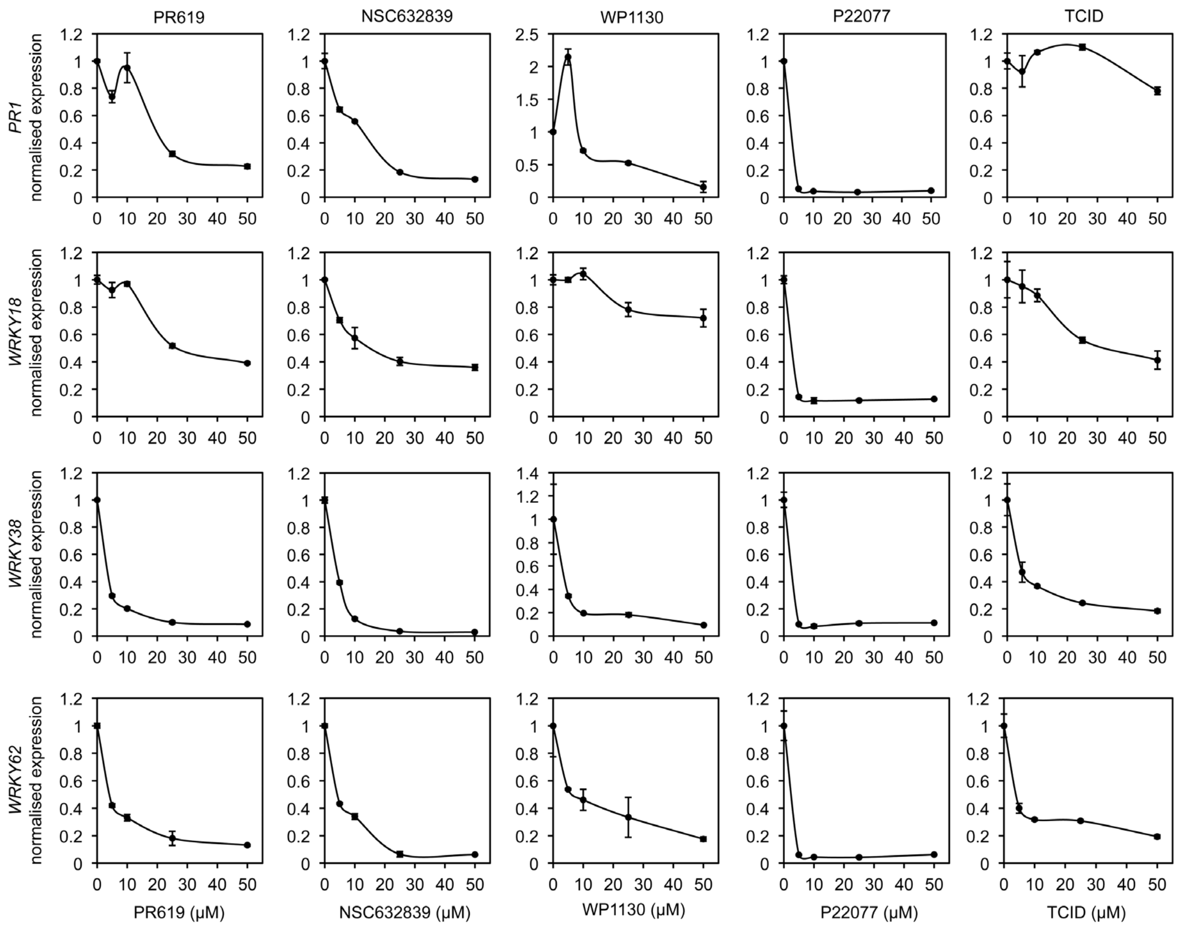


**Figure S3. Related to Figure 4. DUB inhibitors suppress NPR1 target gene expression**

Arabidopsis seedlings were treated with SA in combination with increasing concentrations of the indicated DUB inhibitors. The expression levels of the NPR1 target genes *PR1*, *WRKY18*, *WRKY38* and *WRKY62* were analysed by qPCR and normalised against constitutively expressed *UBQ5*. Data points represent mean relative expression as compared to SA treatment alone ± SD (n=3).

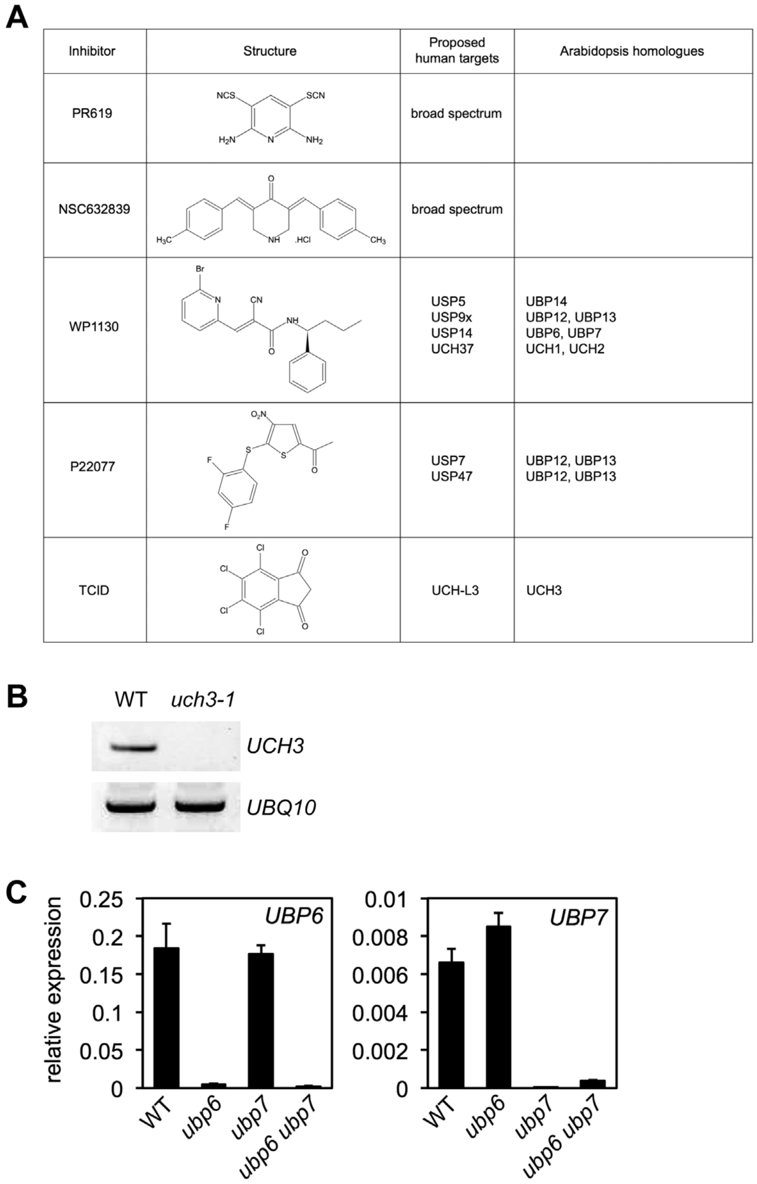


**Figure S4. Related to Figure 5. DUB inhibitor targets in Arabidopsis**

**(A)** Structures and proposed human and Arabidopsis targets of DUB inhibitors used in this study.

**(B)** Expression of *UCH3* was analysed by RT-PCR in the stated genotypes using primers specific to *UCH3* or *UBQ10* as a loading control.

**(C)** The expression of *UBP6* and *UBP7* was analysed by qPCR and normalised relative to constitutively expressed *UBQ5* in plants of the indicated genotypes. Data points represent mean ± SD (n=3).

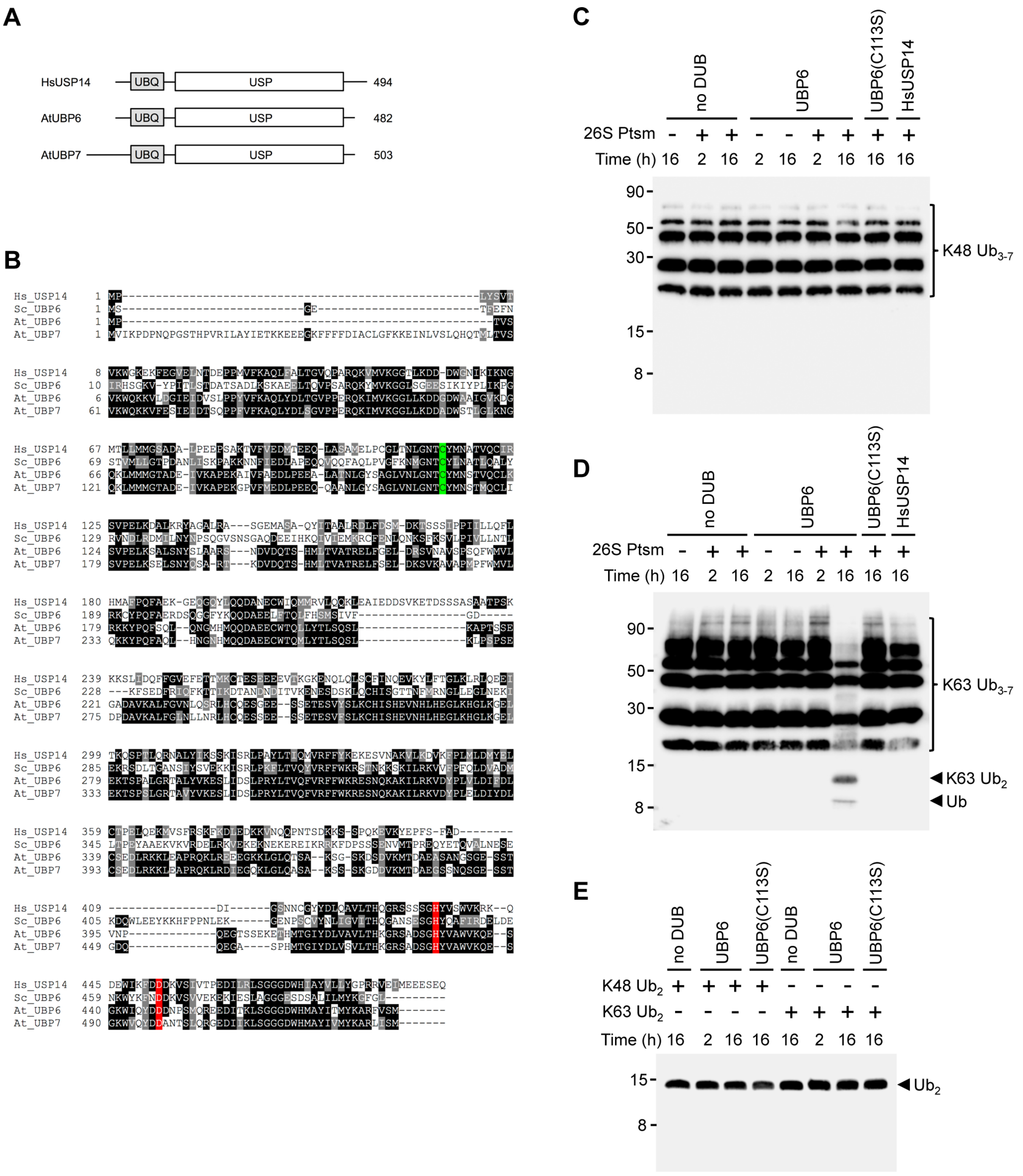


**Figure S5. Related to Figure 6. Structure and activity of UBP6 and UBP7**

**(A)** Domain structures of human USP14, and Arabidopsis UBP6 and UBP7.

**(B)** Sequence alignments of human USP14, *Saccharomyces cerevisiae* UBP6, Arabidopsis UBP6 and Arabidopsis UBP7. The active site Cys residue is highlighted in green while the conserved His and Asp residues making up the catalytic triad are highlighted in red.

**(C)** K48-linked Ub chains of 3-7 in length were incubated with the indicated reaction components for 2 or 16h. Ub species were then detected by SDS-PAGE and immunoblot using antibodies against ubiquitin (P4D1).

**(D)** As in (C) but using K63-linked Ub chains.

**(E)** As in (C) but using K48- or K63-linked di-ubiquitin as a substrate.
